# Unique cuticular morphology of a truly marine insect, *Echinophthirius horridus* von Olfers, 1816 (Phthiraptera: Echinophthiriidae)

**DOI:** 10.1101/2024.10.03.616551

**Authors:** Tucker White, István Mikó

## Abstract

Echinophthiriidae is a family of aquatic lice parasitizing aquatic carnivorans, each member distinguished by their uniquely modified, curved setae. *Echinophthirius horridus* is known to parasitize a wide range of phocid (earless) seals as opposed to exhibiting the more species-specific parasitism of other echinophthiriid lice. In this study, we use a combination of bright field microscopy, confocal laser scanning microscopy (CLSM), and line drawings to provide a detailed description of the general body setae of *E. horridus* and discuss its possible significance as an adaptation to a marine lifestyle.

## Introduction

Earless seals, or phocids, like many marine mammals, face a variety of stressors, from those caused by climate change to those arising from proximity to human activity. These stressors can accumulate and compound one another, leading to possible exponential decrease in overall fitness (Board, 2017). As parts of a dynamic and shifting ocean, marine mammals such as phocid seals will likely benefit from a more holistic approach to their conservation. Legislation such as the Marine Mammal Protection Act of 1972 (Fisheries, 2023) has made significant progress in limiting potential human impact on marine mammals. However, there are many environmental stressors that are less well documented, such as those involving ectoparasitic arthropods and their host interaction dynamics (Rolbiecki et al., 2018).

Echinophthiriidae is a family of sucking lice (Phthiraptera: Anoplura) that are common ectoparasites of aquatic mammals including pinnipeds and the North American river otter (Leonardi and Palma, 2013). While insects like the sea skaters (*Halobates*) are also considered marine, the echinophthiriid lice are considered the only fully aquatic marine insect due to their routine exposure to the deep-water dives and resulting hydrostatic forces they are exposed to by their hosts (Leonardi et al., 2021).

Among echinophthiriid lice, *Echinophthirius horridus* (von Olfers, 1816) is notable for parasitizing multiple species of pinnipeds within the family Phocidae, while other members of Echinophthiriidae are species-specific (Hirzmann et al., 2021). Because of their hematophagous lifestyle, echinophthiriids can spread bloodborne pathogens such as the bacterium *Bartonella henselae* (Morick et al., 2009) and serve as intermediate hosts for endoparasites such as the seal heartworm *Acanthocheilonema spirocauda* (Leidy, 1858), (Ebmer et al., 2022). Due to the wide range of phocid seals parasitized by *E. horridus*, the study of its morphology and host interactions can provide insight into the potential impacts it has across multiple host species.

For ectoparasitic arthropods, the mechanisms by which they attach to their host may involve cuticular specializations, or specific arrangement of cuticular structures, that, in the case of *E. horridus*, allow it to remain attached to their host while exposed to the flow of water. Among lice, cuticular specializations are often host-specific, and many species of lice are only known for parasitizing a single group or even a single species. Phylogenomic sequencing of different species within Echinophthiriidae has shown a consistent pattern of host codivergence with minimal host switching (Leonardi et al., 2019).

Because of the ubiquity of host-specific specialization across lice, the study of the morphology and distribution patterns of cuticular specializations can provide valuable insight into the biology, ecology, and evolutionary history of these parasites. In the case of phocid seals, the study of the cuticular specializations of their lice can illuminate their impact on wild seal populations, and lead to the development of new methods of control or treatment of infestations.

Echinophthiriidae is characterized by its specialized setae that are absent from other anoplurans (Leonardi et al., 2009; Kim, 1986), including highly modified setae covering the dorsal surface of their body (Mehlhorn et al., 2002; Kim, 1971). These specialized setae may be an adaptation to their marine lifestyle, as specimens from the genus *Antarctophthirus* have been observed using their modified setae to hold and coat themselves with the sebum of their host, possibly to serve as a layer of thermal insulation (Mehlhorn et al., 2002).

In this study, we use a combination of bright field microscopy, confocal laser scanning microscopy (CLSM), and line drawings to provide a detailed description of the general body setae of *E. horridus* and discuss their possible significance as an adaptation to a marine lifestyle.

## Materials and Methods

Specimens of the present study (Appendix 1) are deposited at the University of New Hampshire Entomological Collection. Morphological terminology follows Kim (1971) and Beder (1990).

Bright-field images were taken with an Olympus CX-41 compound microscope equipped with a Canon EOS Rebel SLR camera. We also generated 3D datasets using a Nikon A1R-HD confocal laser scanning microscope (CLSM) at the University of New Hampshire Instrumentation Center. To exclude emission from the autofluorescing Canada balsam (embedding media of the specimens) we used two longer wavelength lasers for excitation (639.2nm and 560.3 nm) as opposed to the conventionally used 499nm and 540nm lasers ((Michels and Gorb, 2012; Mikó et al., 2016)) and recorded emission spectra on two channels to distinguish tissue-specific autofluorescences. The channel with longer wavelength emission (less sclerotized structures) was assigned a red pseudocolor and the shorter wavelength channel (more sclerotized regions) was assigned green. CLSM micrograph files were then volume-rendered using FIJI (Schindelin et al., 2012).

The arrangement, size patterns, and external morphology discerned from the bright field photographs and the 3D shape of the CLSM micrographs were then synthesized through the creation of line drawings to combine the mutually exclusive aspects of both imaging techniques. Images were created with 0.15mm and 0.20mm Micron pens.

## Results

### General setae of *E. horridus*

The dorsal surface of the head, thorax, and abdomen and the ventral surface of the thorax and abdomen are covered with setae that are characterized by the presence of a basal setal hump. The hump is a flat-topped projection in the basal region of the dorsal setal surface that is delimited by two distally converging lateral edges (h:Figs 1, 2). The humped seta is curved, bending near its insertion in the body, and thus lies appressed to the body (Figs 1, 2). The dorsal (external) side of the humped seta is convex and the ventral (internal) side is flat. The humped setae can be classified into 3 discrete types based on their length: short (type 1) (Figs 1A, 1D, 2A), medium (type 2) (Figs 1B, 1E, 2B), and long (type 3)(Figs 1C, 1F, 2C). One of the lateral edges of the hump forms a median carina that is bent laterally on type 1 (mr: Fig 2A) and is continuous with the lateral edge in type 2 (Fig 2B) and 3 setae (Fig. 2C). On type 2 setae, two, asymmetrical, converging carinae extend just medially of the distolateral setal edges (cr1: Fig.2B). Multiple dorsal-converging carinae are present on type 3 setae that are, laterally, evenly spaced out from each other (cr1, cr2, cr3: Fig.2C). Similarly to the lateral edge of the hump, the converging carinae are continuous with the lateral edge of the seta.

**Figure 1.**
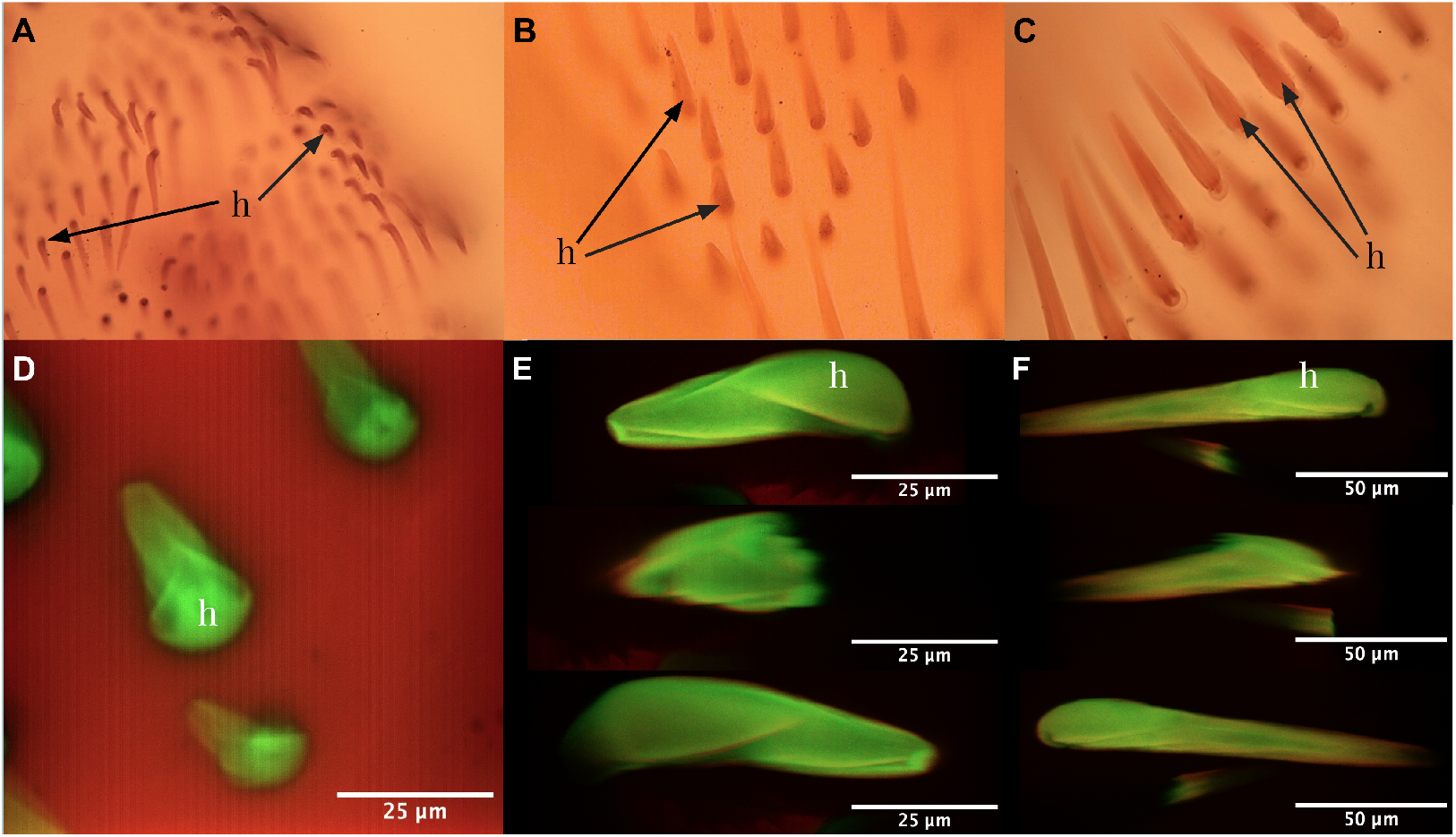
Brght field (A, B, C) and CLSM (D, E, F) images of the setal morphology of *E. horridus* (h=hump).

**Figure 2.**
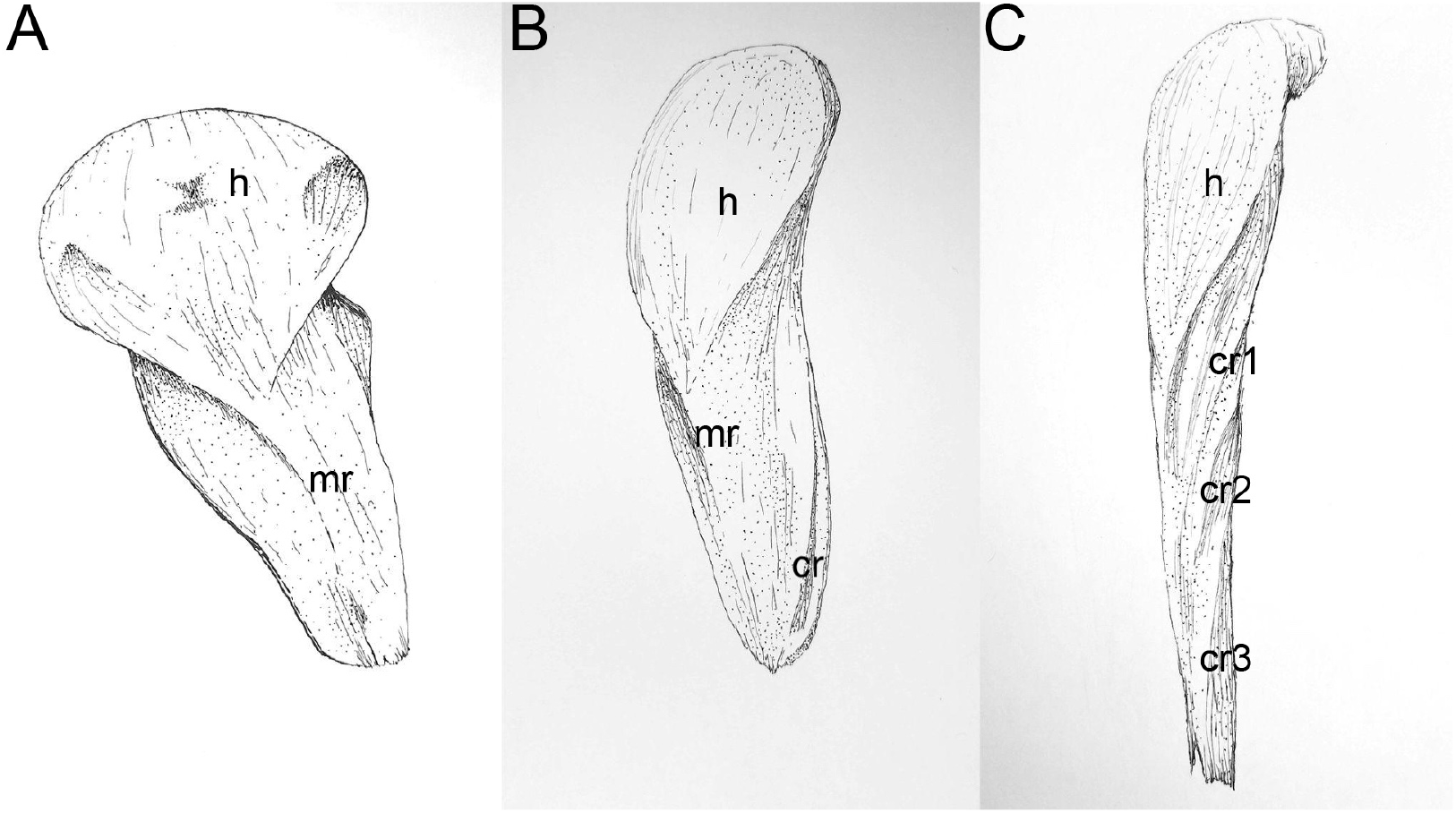
Line drawings of type 1 (A), type 2 (B), and type 3 (C) setae of *E. horridus*.

### Distribution of setal types

Only cranial sclerites possess humped setae on the ventral head region (cs: Fig 3A-D) while the cranial sclerites are devoid of humped setae on the dorsal head region (Figs 3C, D). The cranial sclerites divide the dorsal head region into an hourglass-shaped anteromedian, two anterolateral, and two posterolateral regions. The anteromedian region is exclusively covered with type 1 setae, and the anterolateral and posterolateral regions are covered with type 1 setae anteromedially and with type 2 setae posterolaterally. Three type 3 setae are present along the posterior edges of the posterolateral regions.

**Figure 3.**
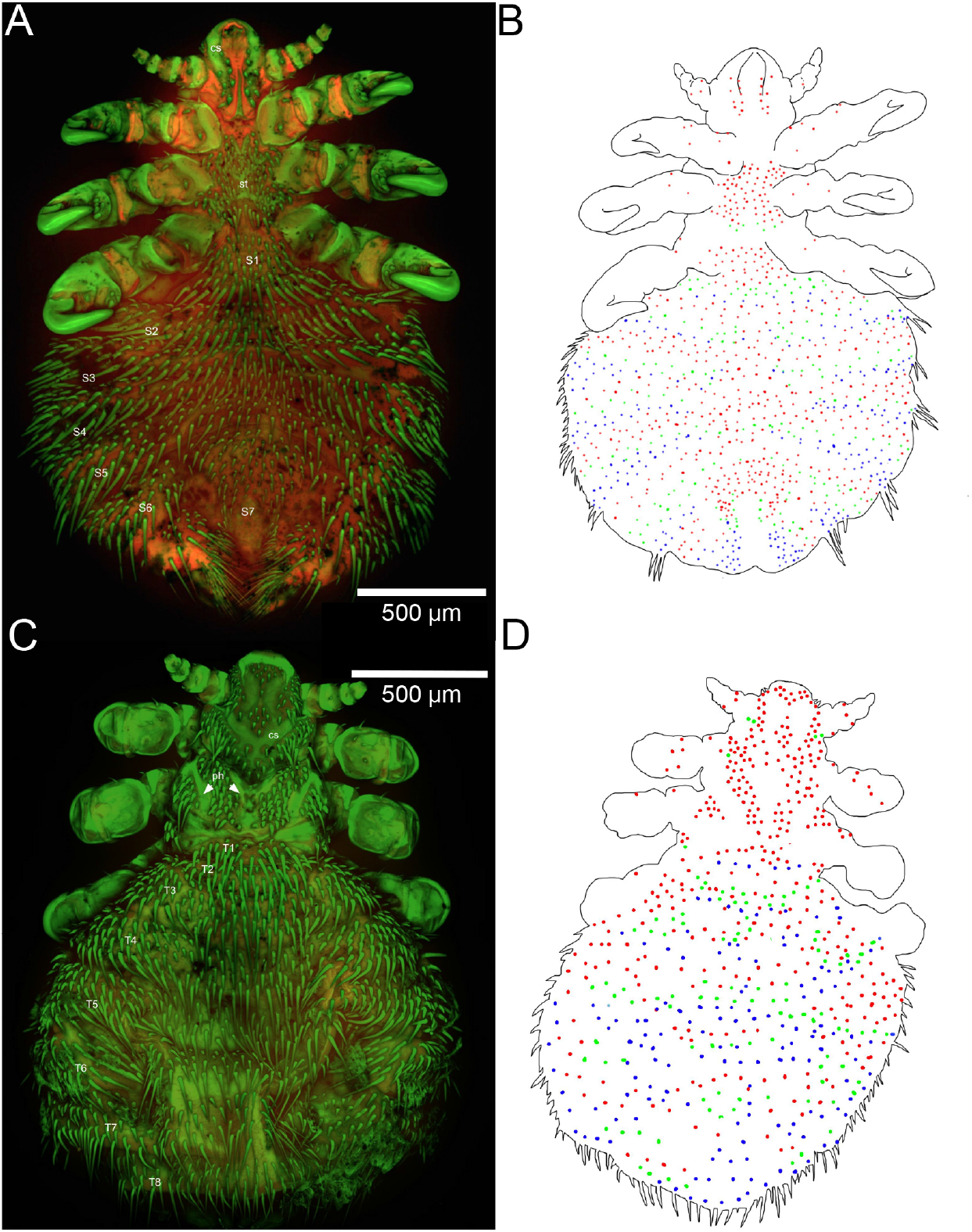
CLSM images of the dorsal (3C) and ventral (3A) sides of *E. horridus* with labelled tergites and sternites respectively. Figures 3B and 3D depict the approximate distribution of the three lengths of setae on both the dorsal (3D) and ventral (3B) sides, using colored dots: Red for type 1, green for type 2, and blue for type 3.

Sclerites of the thorax (phragmata and sternites, ph, st: Figs 3A, C) are glabrous. Conjunctival regions of the dorsal surface of the thorax are covered in mostly type 1 setae with pairs of type 2 setae on the mesothorax and metathorax projecting externally and posteriorly from the sides. The ventral surface contains a uniform spread of type 1 setae, with eight type 2 setae extending along the posterior edge of the metasternum.

One row of type 1 setae extends along the anterior margins, one row of type 2 setae in the middle and one row of Type 3 setae along the posterior margins of the first two abdominal terga. The subsequent terga have multiple setal rows of type 2 setae between the anterior row of type 1 setae and posterior row of type 3 setae. Type 1 setae become less frequent with each additional segment. The setal pattern of abdominal sterna are similar to the 3rd and subsequent abdominal terga except setae are generally shorter in the middle of each tergum. The posterior edge of segment 4 goes from type 3 to type 2 towards the middle, and from segment 4 onwards, the setae closest to the center of each segment decrease in length, culminating in a small bald patch immediately preceding the terminalia on the final abdominal segment.

## Discussion

Scale-like setae with setal hump and the cluster of parallel carinae on type 3 setae are present in all echinopthiriid taxa (Kim, 1986; Smith, 2002; Leonardi et al., 2009), but has not been recorded from any other arthropod taxa (Winterton, 2009; Garm et al., 2013; Richards and Richards, 1979; Larsen, 2003; Cate and Derby, 2001). This unique setal morphology appears to be an adaptation to a fully marine lifestyle, as it is absent from terrestrial lice, including the sister group of Echinophthiriidae, Haematopinidae Kim and Ludwig (1978). Diverse fibrillar attachment mechanisms are well documented across many different insect lineages, including those in underwater environments Chen et al. (2014), and within parasitic arthropods specifically, hook-like structures are a common mode of long-term stability on a host Gorb (2001). The curved setae present on *E. horridus* may have developed to serve a similar mechanical function. Additionally, the grooves on the surface of the curved setae may contribute to the static friction between the setae and the surface of the host seal through increased surface area Gorb et al. (2002). The increased surface area underneath the curved setae might allow for easier retention of the host’s sebum Mehlhorn et al. (2002). Sebum acts as both a thermoregulatory and water-resistant coating in mammalian fur Zouboulis et al. (2003). Coating its body with a layer of its host’s sebum might be an adaptation of *E. horridus* to maintain a constant internal temperature in colder marine environments.

## Acknowledgement

A heartfelt thank you to the staff and students at the UNH Collections, and Mark Townley (UNH Instrumentation Center). This project was funded in part by a grant from The NSF Terrestrial Parasite Tracker Digitization Project. Research reported in this publication was supported through the University of New Hampshire’s Center for Integrated Biomedical and Bioengineering Research (CIBBR) through a grant from the National Institute of General Medical Sciences of the National Institutes of Health under Award Number P20GM113131”.

